# Blood-to-tissue translation in autoimmune disease: paired single-cell evidence from systemic sclerosis

**DOI:** 10.64898/2026.04.18.719421

**Authors:** Niket Rajeevan, Zain Khan

## Abstract

The biology that governs progression and therapeutic response in autoimmune disease is organized in affected tissue, but direct molecular readout of that biology requires invasive biopsy and is rarely repeated during clinical trials or routine care. Using paired blood–skin single-cell RNA-sequencing from a systemic sclerosis (SSc) cohort of 74 individuals (57 patients and 17 matched controls, 192,809 cells across 53 annotated cell states), we show that peripheral blood carries a recoverable projection of tissue-resident molecular state. Across 63 pathways scored in both compartments, 43 same-pathway blood–skin associations reach FDR < 0.05; at cell-type resolution, 212 cross-compartment associations survive residualization for disease status and sex. Per-patient classifiers recover tissue-defined molecular states out of fold with AUCs between 0.62 and 0.79, with the strongest recoveries on fibroblast subtype programs that have no direct circulating analog: fibroblast COMP at 0.79, COCH at 0.75, MYOC2 at 0.74, POSTN at 0.74. Tissue programs route through different blood compartments at different representational levels: fibroblast programs resolve through T-cell, Treg, monocyte and B-cell axes at compositional and distributional levels, while interferon resolves through expression state across multiple cell types. Within SSc alone, a cross-validated partial least squares model learns a shared blood–skin latent axis at *r* = 0.486 (permutation *p* = 0.006); the induced patient ranking recovers tissue-interferon-high patients at 86% precision at the top-20% screening threshold against a 50% base rate. A paired multiview autoencoder, trained on module-level dependency structure under contrastive alignment, paired reconstruction, neighborhood preservation and tissue-target supervision, learns a shared latent geometry in which blood-only projections land in the same tissue-state region as their matched tissue samples and supports recovery of held-out tissue targets above simpler baselines and above two permutation null families. These results map the empirical geometry of cross-compartment inference in autoimmune disease and position peripheral blood as a substrate for tissue-state inference at trial and clinical scale.

## 1 Introduction

The biology that governs progression and therapeutic response in autoimmune disease is concentrated in the affected organ. In SSc the relevant biology lives in the skin and internal organs: dermal fibroblast activation produces the fibrotic remodeling that defines the disease, interferon-stimulated gene programs partition patients into molecular subtypes with distinct prognoses, and vascular endothelial injury precedes late-stage organ damage. Equivalent tissue-localized pathobiology organizes rheumatoid arthritis (synovium), lupus nephritis (kidney), inflammatory bowel disease (intestinal mucosa), and SSc-associated interstitial lung disease (alveolar compartment). The measurements used for patient stratification in clinical trials and for longitudinal monitoring in clinical practice are almost entirely peripheral. The modified Rodnan skin score, the standard primary endpoint in SSc trials, integrates palpation across 17 body sites and does not resolve molecular subtype. Skin punch biopsy resolves subtype but is invasive, incompatible with longitudinal monitoring, and refused at screening by a non-trivial fraction of autoimmune patients. Blood is drawn at every visit.

The gap between what can be measured routinely and what governs clinical behavior is a modeling problem. If peripheral measurements carry a recoverable projection of tissue state, tissue biology becomes computationally accessible from blood without requiring tissue at the point of deployment. The question is whether the projection exists at sufficient fidelity to be useful. We test this using paired blood and skin single-cell RNA-sequencing from Gur et al. [1]. From the 153 individuals profiled in GSE195452, we restricted analysis to 74 with paired blood and skin samples passing quality control (57 SSc, 17 controls), leading to the working dataset that contains 192,809 cells across 53 annotated cell states.

Figure 1 motivates the setup. Blood and skin cells separate cleanly because they are different tissues with different resident populations, and biological organization persists across the compartmental boundary. Immune cells traffic between blood and tissue and acquire transcriptional imprints of the environments they transit; systemic inflammatory mediators reconfigure both compartments in parallel through soluble signaling. Both mechanisms are sufficient to produce cross-compartment alignment, and they predict a bridge that spans beyond any single inflammatory axis.

**Figure 1.**
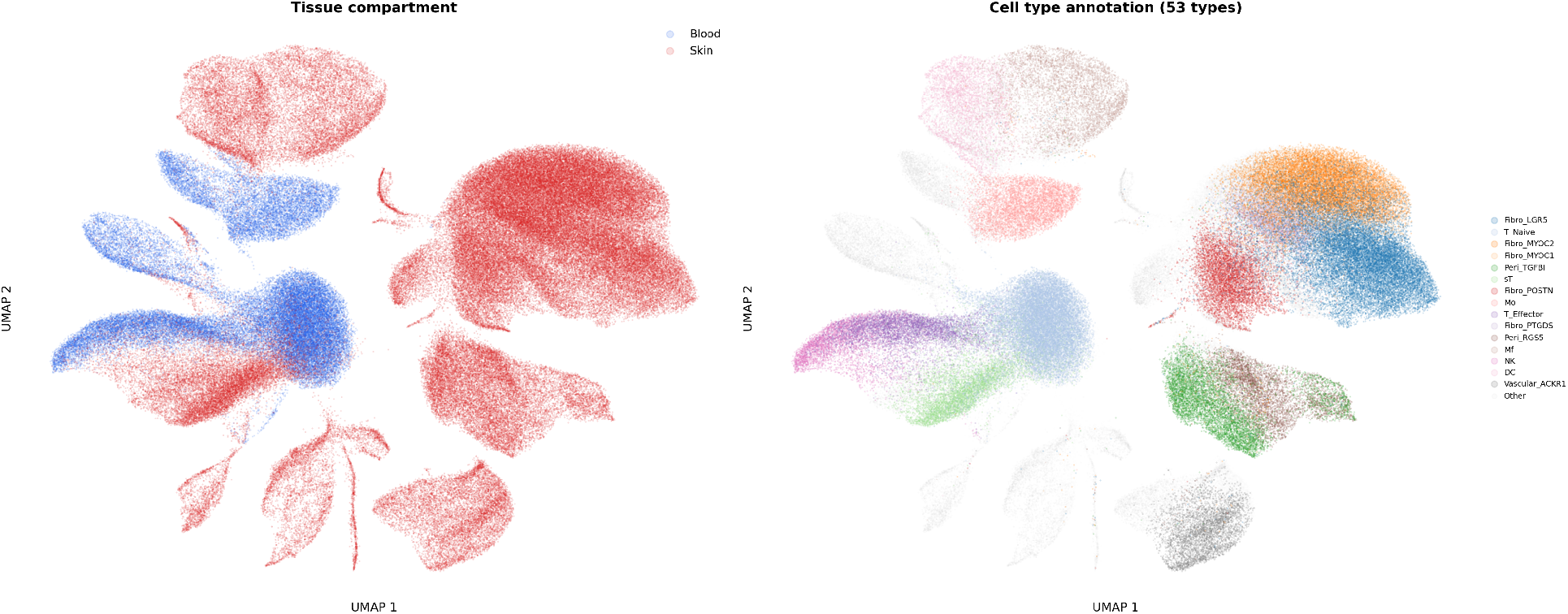
Joint blood–skin single-cell embedding. Left: cells colored by compartment. Right: cells colored by annotation across 53 cell states. Blood and skin occupy distinct regions of the manifold, and biological organization persists across the compartmental boundary: T-cell, monocyte, dendritic-cell and NK populations appear in both compartments with shared transcriptional programs.

This paper establishes the following results for SSc. A blood–skin bridge is detectable at patient level and in shared multivariate structure across 63 pathways scored in both compartments, with 43 same-pathway associations reaching FDR < 0.05 and 212 cross-compartment cell-type associations surviving residualization for disease status and sex. Tissue-defined molecular states are recoverable out of fold from blood alone at AUCs between 0.62 and 0.79, with the strongest recoveries on fibroblast subtype programs that have no direct circulating analogue. Different tissue programs route through different blood cell types at different representational levels, with a wiring pattern that recapitulates known SSc immunopathology from numerical features alone. The bridge persists within disease, produces patient rankings that recover tissue-interferon-high patients at 86% precision at the top-20% screening threshold, and scales: the classifier AUC at *n* = 74 is not saturated, and the single-cell feature interface is compressed at the present cohort size. Finally, a paired autoencoder trained on module-level dependency structure under contrastive alignment and cross-view reconstruction learns a shared blood–tissue latent in which blood-only projections fall in the same region as their matched tissue samples and supports recovery of held-out tissue programs above simpler baselines and above two permutation null families. SSc is the first indication. The target is a cross-indication model whose training interface is paired blood–tissue data at single-cell resolution and whose deployment interface is peripheral blood.

## 2 Data and blood representation

### 2.1 Paired single-cell data

Bulk expression profiles compress thousands of cells into a single vector per compartment and discard compositional and within-cell-state variation. A blood sample measured by scRNA-seq is a distribution over cell states: the abundance of classical versus non-classical monocytes, the fraction of T cells expressing an exhaustion program, the variance of interferon-stimulated gene activity within the NK compartment. Each of these quantities is a potential predictor of tissue state, and none exist in bulk data. When tissue is profiled at the same resolution, fibroblast subtypes and tissue-resident immune populations become individually identifiable targets rather than components of an averaged signal. At 74 paired samples with 192,809 cells across 53 cell states, the Gur cohort is large enough that patient-level distributional features are estimable and cross-compartment associations survive multiple-testing correction across hundreds of feature pairs.

### 2.2 A hierarchical blood representation

Blood carries information about tissue at several biological scales. A patient with expanded circulating monocytes carries compositional evidence of tissue state. A patient whose monocytes are also elevated in interferon-stimulated gene activity carries additional evidence that compositional data alone cannot resolve. A patient whose monocytes display high variance in interferon activity across individual cells, with a long tail of strongly activated cells alongside quiescent ones, carries a third type of evidence that even a per-cell-type average erases. These scales are not redundant, and collapsing them into a single representation discards most of the signal before the modelling begins.

The analytical framework used in sections 3 to 7 interrogates blood at each scale under a common cross-validation protocol and against the same tissue-defined targets. Four representation levels are maintained throughout. Whole-blood pseudobulk is the aggregate expression profile across all blood cells for a given patient, equivalent to what bulk RNA-seq would measure. Cell-type proportions are centered log-ratio transformed to respect the compositional constraint and capture the immune architecture of the sample. Gene-set scores are pathway activity across 55 gene sets — 50 MSigDB Hallmark programs and 5 SSc-specific curated modules for interferon, fibrosis, immune activation, vascular injury and Lgr5-lineage biology — computed both at the whole-blood level and within individual cell types. Single-cell distributional features extract, for each cell type in each patient, the variance of gene-set activity across individual cells, the fraction exceeding a high-activation threshold, and the 90^th^-percentile score; concatenated across eligible cell types and all 55 gene sets, these produce a fixed-length patient vector that preserves the shape of the within-population distribution.

At *n* = 74, the single-cell interface is compressed deliberately. Three robust statistics per cell-type–gene-set pair, a minimum of five cells per patient per cell type, and training-fold median imputation for missing values together define a representation that the present cohort can estimate. Richer distributional features — per-cell embeddings, rare subpopulation frequencies, higher-order co-activation structure — are used by the paired autoencoder in section 8 but are not estimable as stand-alone tabular features at this sample size. The compression is a floor; the scaling behavior in section 7 is consistent with substantial further gain at larger cohort sizes.

### 2.3 Tissue-defined targets

Targets are defined in tissue. Skin gene-set scores and skin cell-state proportions are thresholded at the cohort median to produce binary molecular phenotypes, including tissue-interferon-high, fibrosis-high, immune-activation-high, vascular-high, and fibroblast-subtype-specific labels for COMP, POSTN, MYOC2, Lgr5 and POSTN/PTGDS, defined on the dermal fibroblast subpopulations identified by Gur et al. [1]. These are tissue-resident molecular states rather than clinical severity grades. A fibroblast COMP score reflects the activity of a cartilage-matrix program within a specific dermal fibroblast subtype [2, 3]; circulating COMP protein is elevated in SSc and correlates with mRSS at the bulk level [4], but the identity of the dermal fibroblast subpopulation producing the program is resolved only in tissue. Recovering the subpopulation-level state from blood alone is the core test of every predictive analysis that follows. Blood data does not participate in label construction at any stage.

## 3 Cross-compartment structure

### 3.1 Interferon as framework validation

Interferon is the natural starting point. Type I interferons circulate, plasmacytoid dendritic cells and monocytes in blood are among the strongest responders, and the same interferon-stimulated gene program that marks tissue inflammation is transcriptionally active in the peripheral compartment of high-interferon patients [5, 6]. Blood interferon gene-set score tracks skin interferon score at Spearman *ρ* = 0.593 (*p* = 2.6 × 10^−8^) across all 74 paired individuals (figure 2a). In a mixed cohort any inflammatory axis produces a cross-compartment correlation partly by construction, because patients with disease tend to score higher than controls in both compartments. Within the 57 SSc patients alone the interferon correlation is *ρ* = 0.60 (*p* = 9.3 × 10^−7^) (figure 2b). The variance being explained is the difference between an SSc patient with an interferon-driven skin program and one with quiescent skin, among patients who share the same diagnosis.

**Figure 2.**
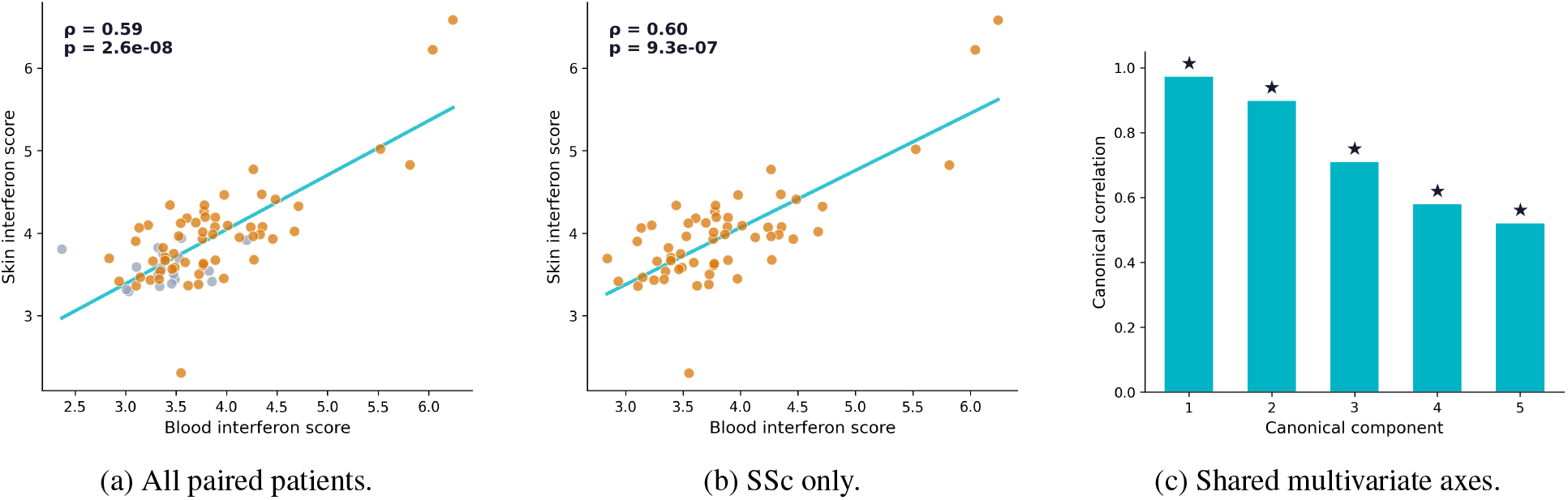
A blood–skin bridge is visible in SSc. (a, b) Blood interferon score tracks skin interferon at patient level in the full cohort (*ρ* = 0.593) and within SSc alone after controls are removed (*ρ* = 0.60). (c) Canonical correlation spectrum across the paired gene-set space. The leading axis (*r* = 0.803, *p* = 0.001) is anchored by the shared interferon program; lower-rank axes carry significant shared signal in proliferative, metabolic and stress-response programs.

### 3.2 Multivariate dependence

Canonical correlation analysis on the full paired gene-set space shows shared structure with dimensionality beyond one inflammatory axis. The leading canonical axis has *r* = 0.803 (permutation *p* = 0.001), anchored by the interferon program in both compartments (figure 2c). The second and third canonical axes carry statistically significant shared signal covering proliferative biology (MYC targets, E2F targets), stress and damage response (p53 pathway, UV response, DNA repair), and metabolic reprogramming (oxidative phosphorylation, glycolysis, fatty acid metabolism). The biological basis is indirect coupling: circulating immune cells traffic through inflamed tissue and acquire a transcriptional imprint of the microenvironment they encounter [7], and systemic mediators reconfigure both compartments in parallel through programs like oxidative phosphorylation and glycolysis that operate in tissue and blood simultaneously [8]. Either mechanism is sufficient to produce cross-compartment alignment on programs that share no single circulating analyte.

Mutual information, which captures nonlinear dependence that CCA as a linear method underestimates, reinforces the scale of this structure. Total observed blood–skin MI across the first five principal components is 2.230, against a permutation null of 0.480 ± 0.168 — a 4.6× excess (*p* = 0.002, 500 permutations). Among the 63 gene sets scored in both compartments, 43 same-pathway blood–skin associations reach significance at FDR < 0.05 after Benjamini–Hochberg correction.

### 3.3 Cell-type coupling

Across all pairwise combinations of blood cell-type features (CLR-transformed proportions and pathway scores) and skin cell-type features, 212 associations reach FDR < 0.05 after residualization for disease status and sex. Residualization matters: without it, many blood–skin correlations survive because SSc patients differ from controls in both compartments, and those associations reflect the dominant disease-versus-health axis rather than within-disease stratification.

After hierarchical clustering (figure 3) the residualized matrix resolves into discrete blocks. Blood monocyte and macrophage features cluster with skin fibroblast populations, consistent with monocyte recruitment to SSc skin [9]. Blood dendritic-cell features cluster with skin immune activation scores [10]. Blood T-cell features, particularly Treg and effector subsets, connect to skin stromal and vascular programs [11, 12]. These associations run between circulating immune populations and tissue-resident programs of different cellular identity: a blood monocyte feature predicting a skin fibroblast program is consistent with the imprint that a fibroblast-active tissue environment leaves on the circulating immune system. The fibroblast recovery results in section 4 depend on exactly these indirect routes.

**Figure 3.**
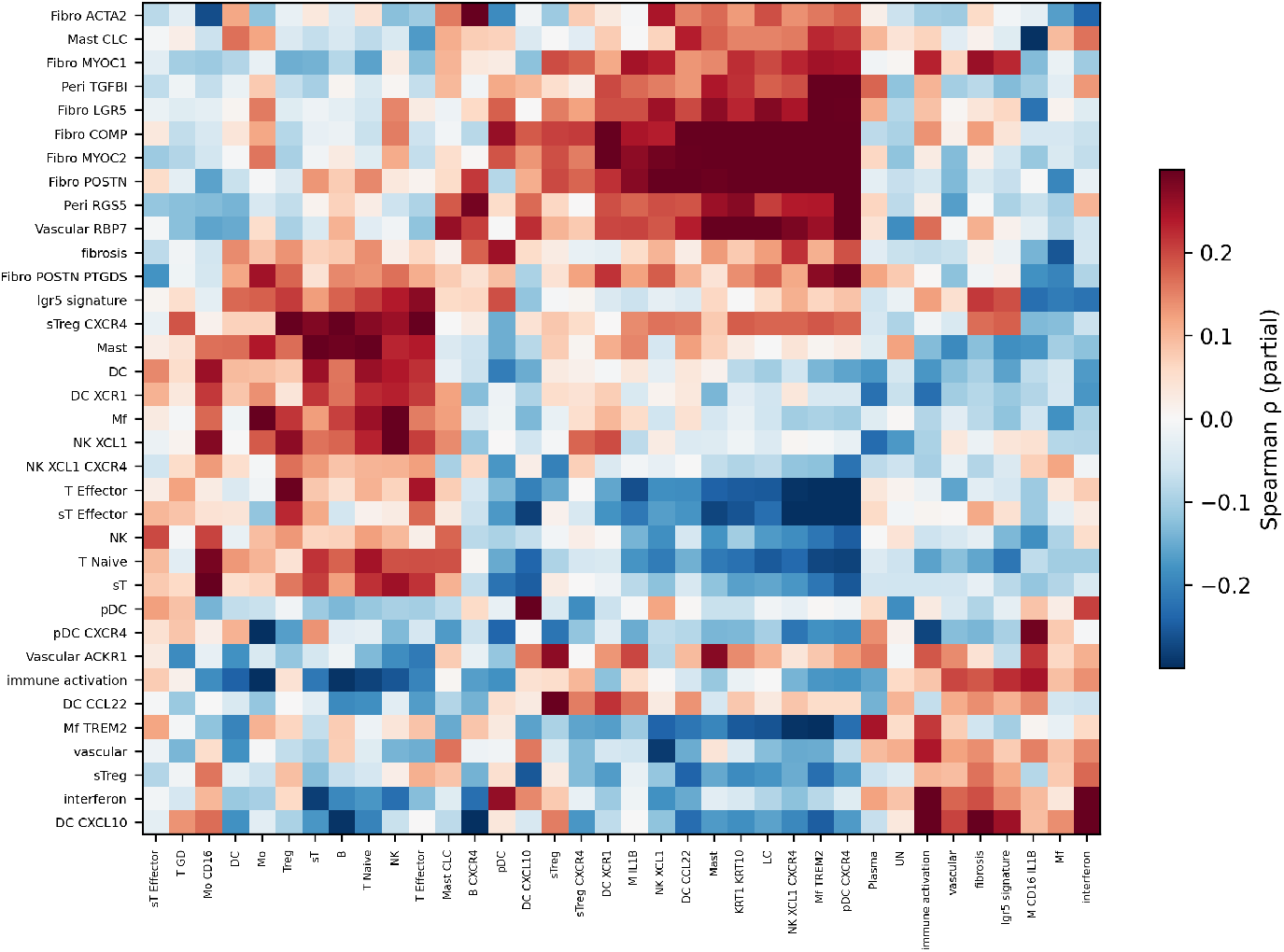
Residualized cross-compartment association map. Rows: blood features. Columns: skin features. Hierarchical clustering resolves the matrix into discrete blocks. Blood monocyte and macrophage features link to skin fibroblast programs; blood dendritic-cell features link to skin immune activation; blood T-cell features link to skin stromal and vascular states.

## 4 Recovery of tissue-defined states from blood

The associations in section 3 describe statistical coupling at the population level. The predictive question is harder: given only blood from a held-out patient, can a model recover which tissue-defined molecular state that patient carries?

### Protocol

Patients are labeled tissue-high or tissue-low by median-splitting their skin gene-set scores or cell-state proportions. The classifier predicts the tissue-defined label of a held-out patient from blood features alone. Cross-validation is Repeated class-wise stratified K-Fold, 5 folds × 2 repeats, with all feature construction (HVG selection, StandardScaler fitting, PCA, distributional z-score thresholds) performed strictly inside training folds. Out-of-fold predicted probabilities are averaged across repeats per patient. The classifier is a random forest (200 trees, max depth 4, minimum leaf size 3) regularized for *n* = 74. Every tissue target is evaluated at every blood representation level (pseudobulk PCA, proportions, gene-set means, single-cell distributional features, and combined) under identical folds and classifiers.

### 4.1 Per-target recovery

Figure 4a shows the best out-of-fold AUC achieved by any blood representation for each tissue target. AUCs range from 0.62 to 0.79. The strongest recoveries are on tissue programs with no direct circulating analogue. Fibroblast COMP reaches 0.79 from blood cell-type proportions. COMP is a cartilage-matrix gene expressed by a specific dermal fibroblast subtype; while serum COMP is elevated in SSc as a bulk fibrosis proxy [13], fibroblasts themselves do not circulate, and the identity of the fibroblast subpopulation producing the program is resolved only in tissue. What proportions capture is the compositional signature that a COMP-active tissue environment leaves on peripheral immune architecture. Fibroblast COCH (0.752) and fibroblast MYOC2 (0.742) follow the same pattern, both recovered from proportions. Fibroblast POSTN reaches 0.737 from single-cell distributional features rather than proportions, a distinction examined in section 4.2. DC-state reaches 0.739 from whole-blood pseudobulk; plasma cell programs reach 0.717 from proportions, as expected for a population that circulates as plasmablasts before tissue homing [14]. Tissue interferon reaches 0.709 from gene-set mean scores — the aggregate pathway score across all blood cells — because interferon elevates the same transcriptional program across multiple blood cell types simultaneously [6] and decomposing it into per-cell-type structure adds marginal information.

**Figure 4.**
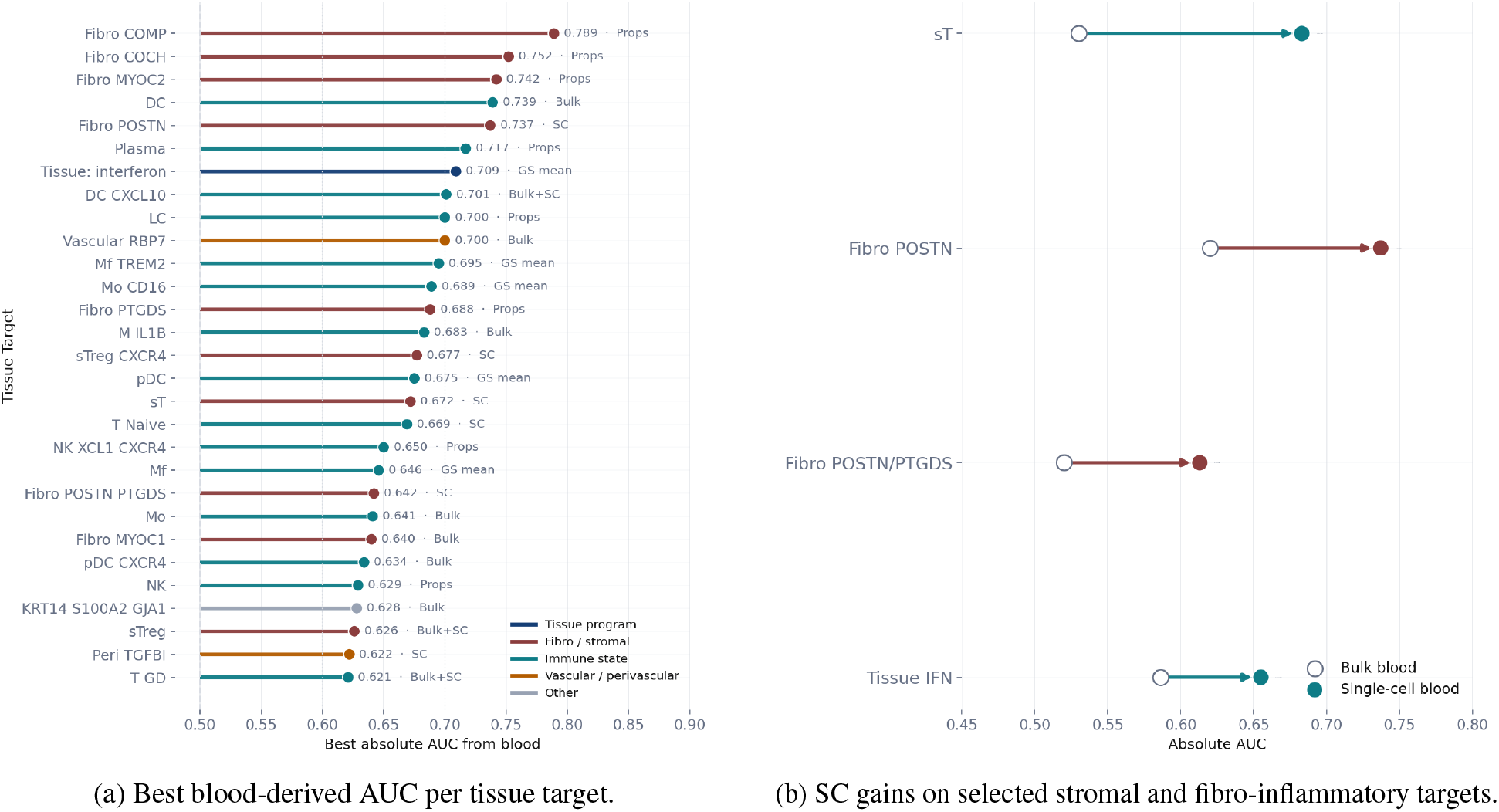
Blood-derived classifiers recover tissue-defined molecular states. (a) Best out-of-fold AUC per tissue target, colored by the blood representation that achieved it. AUCs range from 0.62 to 0.79. (b) For selected targets, single-cell distributional features (filled) versus whole-blood pseudobulk (open). The largest SC gains appear on fine-grained stromal and fibro-inflammatory targets where within-cell-type heterogeneity carries discriminative information that pseudobulk averaging erases.

Further along the recovery spectrum, immune activation (0.695 from gene-set scores), Mf TREM2 (0.659), monocyte CD16 (0.656), stromal T cells (0.671 from SC features), fibroblast POSTN/PTGDS (0.661) and T GD (0.621) recover above chance with weaker discrimination. These targets couple to blood through distributional rather than compositional mechanisms and are the targets most likely to sharpen as paired sample sizes grow (section 7).

### 4.2 Resolution dependence

Which blood representation best recovers which tissue target follows a biologically interpretable gradient (figure 4b). Coarse whole-compartment programs — interferon, broad immune activation, DC state — are already well recovered from pseudobulk or gene-set mean scores. These are programs where blood–tissue coupling operates through a coordinated transcriptional shift across many cell types at once.

Fine-grained stromal and fibro-inflammatory targets show directional gains under SC features relative to pseudobulk. Fibroblast POSTN, fibroblast POSTN/PTGDS and stromal T-cell programs couple to blood through a distributed mechanism: a fibroblast POSTN-active tissue microenvironment does not elevate one gene module uniformly across the blood, but manifests in distributional shifts within specific cell compartments for example tail behavior in NK cells or heterogeneity of Treg engagement. These signals exist in the shape of within-cell-type distributions; pseudobulk, by averaging, destroys them. The SC advantage on these targets confirms that within-cell-type heterogeneity carries recoverable tissue information.

### 4.3 Permutation controls

Blood features are high-dimensional and internally correlated. A flexible classifier could in principle exploit within-blood correlations to produce above-chance AUCs on tissue labels whose median split also partitions the cohort into two groups of similar size. The negative control tests this directly. Each patient’s blood features are preserved; the tissue label assigned to each patient is drawn randomly from the cohort.

A formal permutation test (500 permutations) on the interferon target produces null distributions centered on AUC ≈ 0.50 with standard deviation 0.07–0.09, consistent with the variance of the AUC estimator at *n* = 74. SC distributional features achieve permutation *p* = 0.040 (observed AUC 0.655, null 0.505 ± 0.084). Whole-blood pseudobulk at observed AUC 0.564 does not reach significance (*p* = 0.214, null 0.503 ± 0.071); cell-type proportions at 0.509 are indistinguishable from the null (*p* = 0.481). Under shuffled labels predictive structure collapses to chance (figure 5a). The predicted probability distributions in figure 5b,c show that even on interferon, where an aggregate gene-set score achieves the highest AUC, decomposing blood into per-cell-type distributional structure sharpens the posterior separation between tissue-high and tissue-low patients.

**Figure 5.**
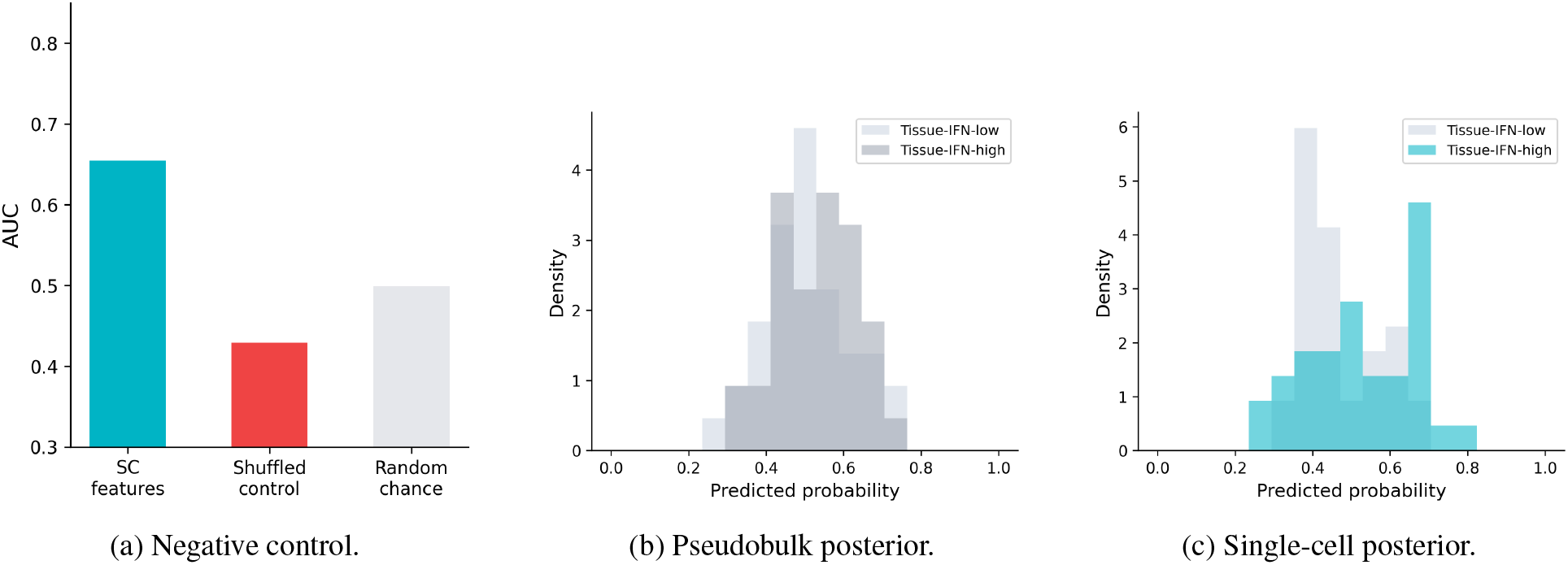
Permutation control and posterior calibration on the tissue-interferon target. (a) Shuffled patient labels collapse predictive structure to chance. (b, c) Predicted probability distributions for tissue-IFN-high and tissue-IFN-low patients under whole-blood pseudobulk (b) versus single-cell distributional features (c). SC features produce sharper posterior separation on a target where aggregate gene-set scores achieve the highest AUC.

## 5 Cell-type routing

The per-target recoveries in section 4 report the best AUC across all blood representations pooled. A complementary analysis asks which specific blood cell type and which information channel carry each tissue target. Figure 6 trains a separate classifier restricted to a single blood cell type and a single channel (proportion, SC distributional, pseudobulk, gene-set score) for each tissue target. Each bubble is a cell-type–channel–target combination achieving AUC ≥ 0.60 out of fold. The classifier receives only numerical features, so the wiring pattern is an empirical finding rather than a prior.

**Figure 6.**
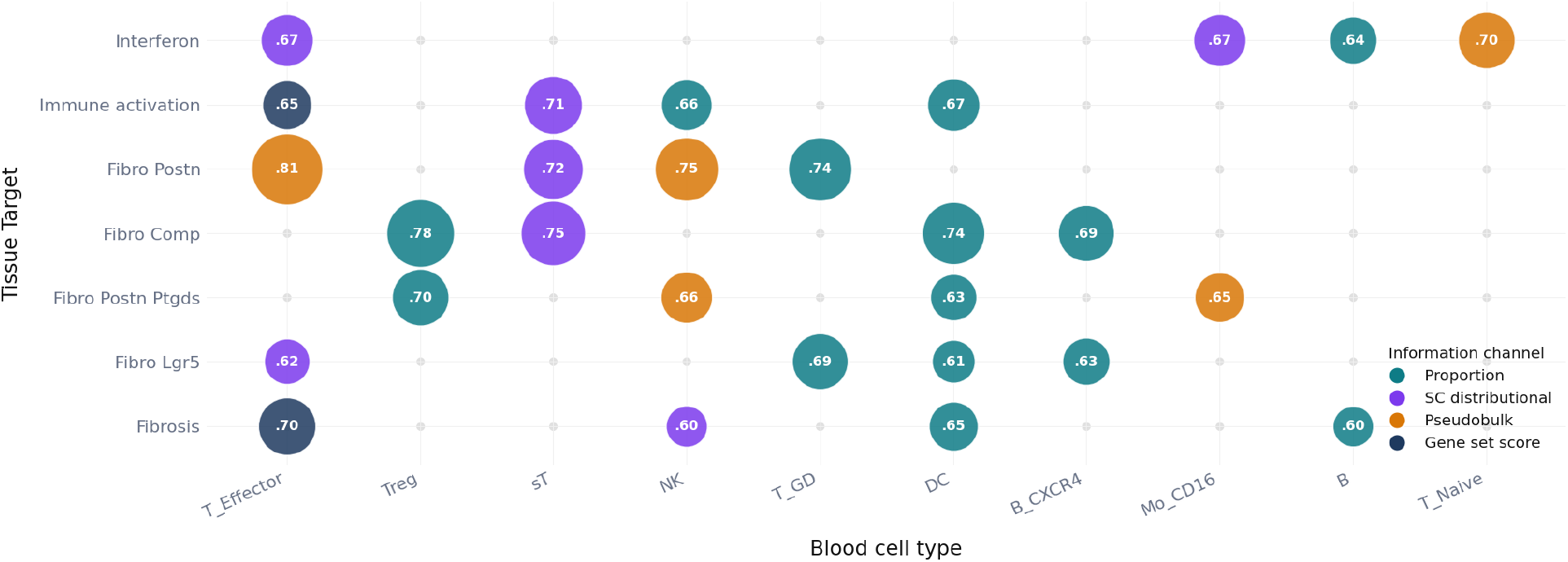
Per-cell-type recovery of tissue targets across four blood information channels. Each bubble: one blood-cell-type–channel–target combination at AUC ≥ 0.60 out of fold; size scales with AUC. Fibroblast programs route through T-cell and Treg proportions with secondary monocyte and B-cell axes for COMP. Interferon routes through pseudobulk and gene-set scores across multiple cell types. The wiring is target-specific and recapitulates known SSc immunopathology from numerical features alone.

### Fibroblast POSTN routes through the T-cell compartment

T-effector proportions alone recover fibroblast POSTN at 0.81, the strongest single cell-type–target edge in the map. Treg proportions reach 0.72, skin-tropic T cells 0.72, NK cells 0.75 and gamma-delta T cells 0.74. The wiring has a mechanistic basis in the SSc literature. Peripheral blood effector CD8^+^ T cells in SSc overproduce IL-13 at levels that correlate with skin fibrosis extent [15], and IL-13-producing CD8^+^ T cells are the predominant early infiltrate in SSc skin where they induce a pro-fibrotic phenotype in dermal fibroblasts [11]. CD4^+^ cytotoxic T lymphocytes in SSc drive endothelial apoptosis and fibroblast activation through cytokine secretion [16]. Th17/Treg imbalance in peripheral blood tracks fibrosis severity [17, 18], and Tregs that enter SSc skin undergo tissue-specific conversion into TH2-like cells producing IL-4 and IL-13, a plasticity driven by local IL-33 that is absent in the blood Treg compartment of the same patients [12]. A patient with active POSTN-expressing fibroblasts has a tissue microenvironment that selectively recruits and converts circulating T-cell populations; the recruitment reshapes blood T-cell composition, and the classifier recovers the reshaped composition from proportions.

### Fibroblast COMP recruits monocyte and B-cell axes

Fibroblast COMP shares the Treg axis with POSTN (0.78 from Treg proportions) but recruits B_CXCR4 proportions at 0.74 and Mo_CD16 proportions at 0.69. The monocyte axis reflects a distinct arm of SSc fibrosis: M2-polarized macrophages differentiate from recruited circulating monocytes and drive fibroblast-to-myofibroblast transition through TGF-*β* and PDGF secretion [19], with elevated soluble CD163 found in blood and skin of early SSc patients [20]. The B_CXCR4 axis is consistent with broader dysregulation of the SSc B-cell compartment, including Tfh expansion, where B-cell-derived IL-6 contributes to the pro-fibrotic cytokine milieu [21]. A further distinction: the skin-tropic T-cell signal for COMP comes through SC distributional features (0.75) rather than proportions, indicating that for COMP the discriminative information in the sT compartment lives in the shape of the within-population activation distribution rather than in population abundance.

### Interferon routes through expression state

Interferon resolves through qualitatively different wiring. Mo_CD16 pseudobulk at 0.67, B-cell pseudobulk at 0.64, T-naive gene-set scores at 0.70 and T-effector proportions at 0.67 are the dominant carriers. The information channels shift from compositional and distributional (proportions and SC) to expression state (pseudobulk and gene-set scores). Type I interferons are produced in large part by plasmacytoid dendritic cells, whose depletion in mouse models of SSc prevents and reverses fibrosis [10], and the resulting interferon-stimulated gene module activates coherently across monocytes, B cells, T cells and NK cells simultaneously [6]. The tissue-interferon signal is carried in what blood cells are transcribing rather than in which blood cells are present, which is why expression-state features outperform compositional features on this target while the reverse holds for fibroblast programs.

### Distributed landscape

No single blood cell type is sufficient. T-effector features recover POSTN at 0.81 but carry no signal for the fibrosis axis. Mo_CD16 carries the interferon pseudobulk signal and the COMP proportional signal while contributing nothing to immune activation; B_CXCR4 appears for COMP and fibrosis but is absent from POSTN and interferon. The columns of figure 6 do not generalize across rows, and the rows do not generalize across columns. The tissue-state landscape of SSc is distributed across cell types and representational levels, with target-specific routing that reflects the distinct pathobiological mechanisms through which each tissue program couples to circulating immunity. The paired autoencoder in section 8 models this distribution directly.

## 6 Within-disease structure and trial enrichment

### 6.1 Joint pathway recovery within SSc

The recovery analyses in section 4 operate on one tissue target at a time and collapse a continuous tissue score to a binary label. A cross-validated partial least squares (PLS) model fit on blood gene-set scores to reconstruct skin gene-set scores, using only the 57 SSc patients and with sex residualization performed inside each training fold, learns a shared latent axis that maximally co-varies between blood and skin across the entire 55-pathway space. The leading PLS axis yields *r* = 0.486 (permutation *p* = 0.006, 500 permutations) on held-out SSc patients. A single latent dimension of blood captures coordinated variation in interferon, fibrotic, metabolic, proliferative and stress-response programs in skin simultaneously, evaluated out of fold and within disease.

The patients separated along this axis are SSc patients with active, multi-program tissue involvement versus SSc patients with comparatively quiescent skin — a distinction that integrates across inflammatory, fibrotic and metabolic axes simultaneously and that currently requires biopsy to make. The tissue-state information in blood has shared low-dimensional structure: multiple tissue programs co-vary with multiple blood programs in a way that a single latent axis already captures at *r* = 0.486 within disease.

### 6.2 Enrichment precision

We evaluated a predefined panel of 14 within-SSc tissue targets spanning the main biological axes of the skin compartment rather than selecting only targets that ranked well post hoc. These 14 targets capture disease-relevant heterogeneity that is biologically meaningful inside the SSc subset, and were interferon, fibrosis, immune activation, vascular, LGR5 signature, fibroblast LGR5 fraction, fibroblast POSTN fraction, T-cell infiltration, myeloid infiltration, stromal T cells, sTreg CXCR4, fibroblast MYOC2, fibroblast POSTN, and fibroblast POSTN/PTGDS. This gives a broad but still interpretable screen of the tissue-program landscape without collapsing the analysis onto a single favored endpoint. At the top-20% ranking cutoff, using the best blood representation for each target, 10/14 targets clear 60% precision, 9/14 clear 70%, and 6/14 clear 80% within SSc.

Within-disease classifiers (SC distributional features, trained within SSc only with tissue targets re-thresholded at the SSc-only median) produce out-of-fold patient rankings that are directly usable for trial enrichment (figure 7).

**Figure 7.**
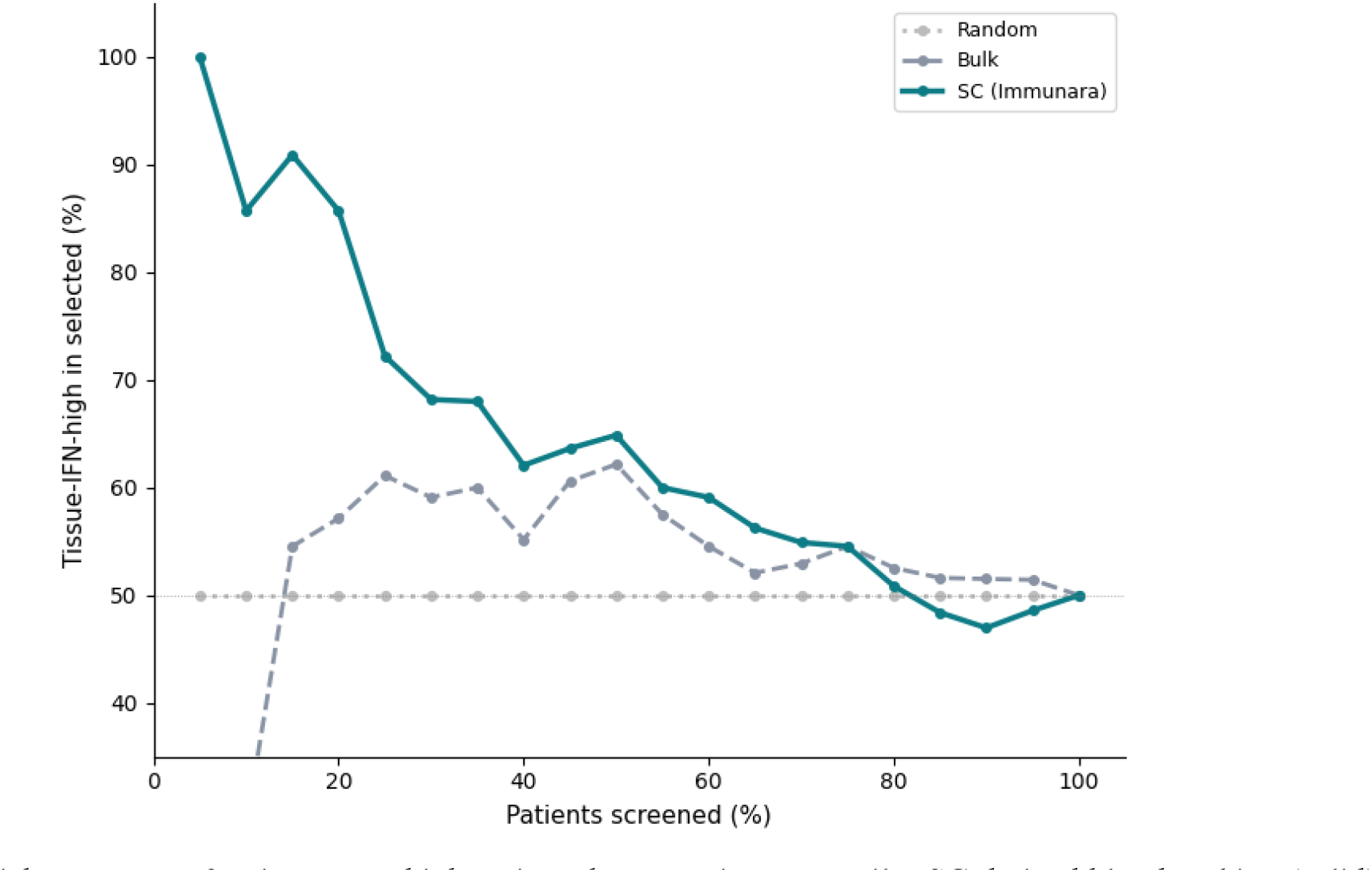
Enrichment curve for tissue-IFN-high patients by screening percentile. SC-derived blood ranking (solid) versus whole-blood pseudobulk (dashed) versus random (dotted). At the top 20%, SC ranking selects tissue-IFN-high patients at 86% precision against a 50% base rate; at top 15%, 91%.

**Figure 8.**
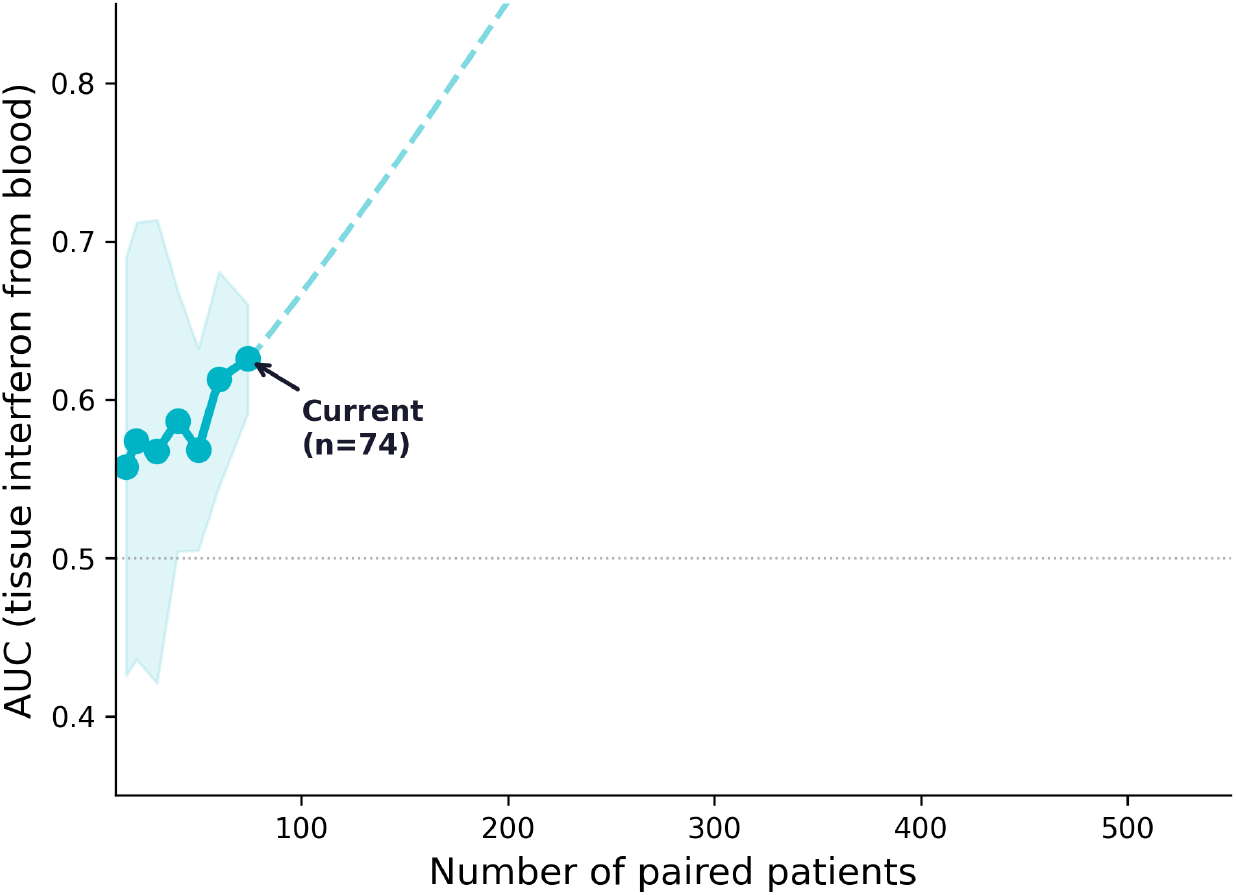
Scaling of blood-to-tissue recovery on the interferon target. AUC against paired cohort size under stratified subsampling (20 resamples per *n*, error bars ± 1 SD). The SC representation improves with sample size; the curve is not saturated at *n* = 74.

**Figure 9.**
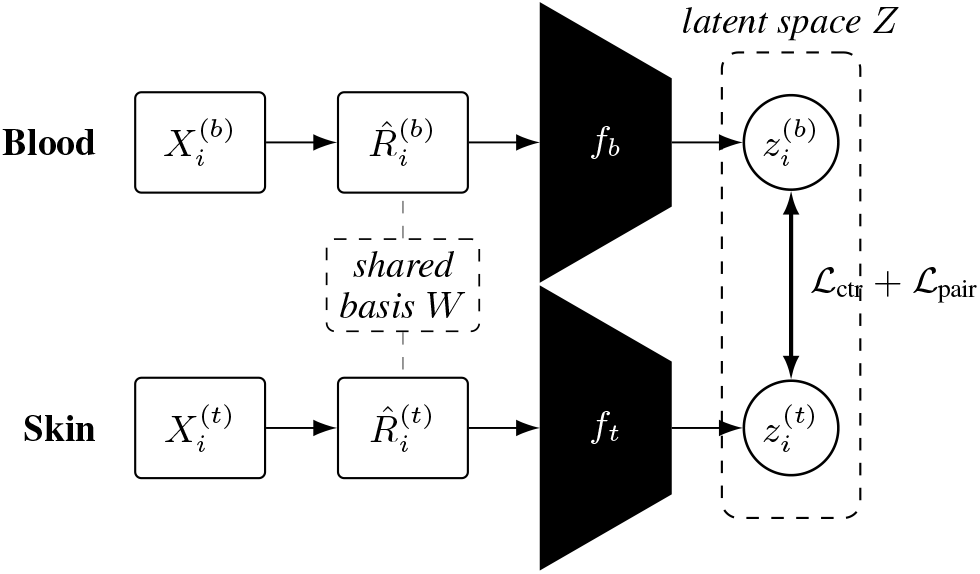
Paired autoencoder architecture. Paired blood and skin single-cell matrices 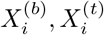 are compressed through a fold-safe shared gene-module basis *W* into patient-level module-by-module shrinkage-Spearman dependency matrices 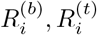, vectorized, and passed through compartment-specific MLP encoders *f*_*b*_, *f*_*t*_ into a shared latent space *Z*. Alignment losses ℒ_ctr_ (symmetric InfoNCE contrastive) and ℒ_pair_ (cosine pull on matched pairs) drive 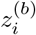 and 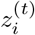 for the same patient together while separating mismatched cross-patient pairs. Auxiliary terms preserve dependency structure under self- and cross-reconstruction (*L*_rec_), preserve local patient neighbourhoods in *Z* (ℒ_nbr_), prevent latent collapse (ℒ_var_), and support continuous tissue-target readout from both compartments (ℒ_breg_, ℒ_treg_).

At the top 20%, 86% of selected patients are truly tissue-IFN-high against a base rate of 50%; at top 15%, precision reaches 91%. Whole-blood pseudobulk under the same cross-validation protocol reaches approximately 57% at top 20%, while random selection yields 50% by definition. The same ranking infrastructure generalizes across tissue targets: within SSc, blood-derived rankings recover 73% of stromal T-cell-high patients at top 20%, 82% of sTreg CXCR4-high patients, 82% of fibroblast MYOC2-high patients, and 88% of fibroblast POSTN-high patients at top 30%. Together these targets span immune infiltration, regulatory T-cell biology, and fibroblast remodeling, showing that the ranking framework is not confined to a single endpoint but operates across a broader tissue-program landscape.

### 6.3 Trial-economic consequences

An SSc Phase 2 trial targeting interferon-driven disease currently faces two options. Enroll unselected patients, accepting that a substantial fraction of the cohort carries quiescent tissue on the interferon axis and will dilute the treatment signal — prior work estimates tissue-IFN-high prevalence at roughly 40–60% across SSc cohorts [24]. Or require skin punch biopsy for molecular stratification, which adds screening attrition and extends the screening timeline. In SSc specifically, biopsy is complicated by skin fragility and delayed wound healing in fibrotic tissue [25]. Biopsy-requiring oncology trials show median screening durations roughly twice those of non-biopsy trials [26]; analogous friction applies when tissue stratification is placed upstream of randomization in autoimmune indications.

Beyond stratification, the blood-derived scores open a second operational surface. The mRSS requires 52 weeks to read because it measures a slow macroscopic consequence of the underlying molecular process. Inter-observer variability is ± 4.6 units [27], and the score has a well-documented tendency to improve on placebo across Phase 2 programmes, contributing to trials in which active arms failed to separate from a declining control [28]. The latent blood–tissue axis learned by PLS [29] tracks coordinated variation in tissue interferon, fibrotic, metabolic and proliferative programs from blood. If the axis is treatment-responsive — a question the present cross-sectional cohort cannot answer — a blood-derived readout could in principle support earlier adaptive decisions than the week-52 mRSS endpoint. Formal qualification as a surrogate requires prospective longitudinal data on treated patients; the current work establishes the axis, not its treatment-responsiveness.

Skin is the most accessible tissue in the autoimmune portfolio: a 4 mm punch biopsy under local anesthetic, performed in an outpatient setting. The closest equivalent in rheumatoid arthritis, ultrasound-guided synovial biopsy, is well-tolerated but requires specialist rheumatology infrastructure and imaging [30]. In lupus nephritis, stratification by tissue requires renal biopsy, which carries bleeding risk and typically mandates overnight observation. The alveolar compartment in SSc-associated interstitial lung disease is accessible only through bronchoscopy or surgical lung biopsy. If biopsy-based stratification already introduces meaningful operational friction even for skin, the case for blood-based tissue-state inference scales with the invasiveness and infrastructure cost of the alternative across the broader autoimmune portfolio.

## 7 Scaling behavior

The scaling analysis asks how recovery changes as the paired cohort grows. At each sample size *n*∈ {15, 20, 30, 40, 50, 60, 74}, patients are subsampled without replacement (stratified to preserve the class ratio) and the full cross-validated pipeline for SC distributional feature construction and random forest training is re-run from scratch. Twenty independent resamples are drawn at each *n* to estimate the mean and variance of AUC. The interferon target is used as the demonstration case because it has the strongest prior biological expectation of cross-compartment signal; fibroblast subtype targets achieve higher absolute AUCs at *n* = 74 (figure 4a), and the qualitative trajectory applies across the target set.

At *n* = 15, AUC fluctuates near chance (≈0.55) with high variance across resamples. At *n* = 74, SC features reach their reported AUC of ≈0.66 on this target. The trajectory from *n* = 50 to *n* = 74 shows no sign of flattening, and the SC curve is still steepening at the largest available sample size.

Two mechanisms drive the continued improvement and act on different parts of the system. The first is classical statistical power: more patients produce tighter estimates of existing feature–target associations and stabilize the classifier’s decision boundary. This mechanism plateaus once the existing feature set is fully exploited. The second is specific to the single-cell interface and has no analogue in bulk. At *n* = 74 the SC features are compressed into three robust statistics per cell-type–gene-set pair because richer distributional features — co-activation structure across gene-set pairs, rare subpopulation frequencies, higher-order moments of the per-cell activation distribution — cannot be estimated reliably from the number of cells per patient available at this cohort size. Many cell-type–patient combinations have fewer than 100 cells, some fewer than 20. At *n* = 200–500, features that are currently too sparse to estimate become statistically usable, and the SC representation widens: new features that do not exist in the current model become estimable and carry tissue information that the present three-statistic compression leaves on the table. The scaling curve at *n* = 74 reflects only the first mechanism and is therefore a lower bound on the SC scaling trajectory.

## 8 Paired autoencoder

We start this section by reinstating the key high-level problem. For each patient we have two paired measurements: a blood single-cell RNA-seq sample and a skin single-cell RNA-seq sample. The scientific question is whether the blood sample contains enough information to recover something about the patient’s tissue state. The ordinary machine learning analyses in sections 3 to 6 answer this using hand-engineered features and target-specific classifiers. The paired autoencoder asks a more ambitious question: can we construct a *shared latent representation* of patient state such that blood and tissue are two views of the same hidden disease geometry?

### 8.1 Fold-safe gene-module basis

For each patient *i* and compartment *c* ∈ {*b, t*}, the input is a sparse single-cell expression matrix 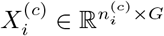, with rows indexing cells and columns indexing genes. The raw matrix is not fed directly into the patient-level model; the number of cells 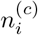 varies across patients and is small for many cell-type–patient combinations, and the gene-level representation is redundant with respect to the biological programs that distinguish patients.

Inside each training fold only, a shared gene-module basis is constructed. From the training patients’ cells (up to a cap of 20,000 cells for tractability), the top *H* = 256 highly variable genes are selected, each gene is standardized to training-fold mean and variance, the *H* × *H* gene–gene Spearman correlation *C* is computed, converted to distance *D*_*gg′*_ = 1 − *C*_*gg′*_, and average-linkage hierarchical clustering is applied with a cut at *M* = 32 modules. The resulting module assignment map *m*(*g*) ∈ {1, …, *M}* is a function of the training fold only, so the basis is fold-safe: no test patient contributes to basis construction. Applied across all patients in the fold, this converts each cell from a *G*-dimensional gene vector to an *M*-dimensional module-score vector.

### 8.2 Patient-level dependency representation

For patient *i* and compartment *c*, per-cell module scores are computed by averaging the standardized expression of the genes assigned to each module,

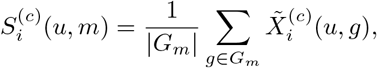

yielding a cells-by-modules matrix 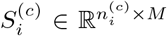. The patient-level object is the module-by-module Spearman correlation across the cells in the sample,

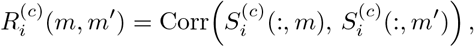

shrunk toward the identity to stabilize at finite cell counts:

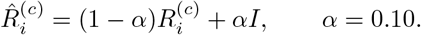

Vectorizing the strict upper triangle gives the model’s input vector,

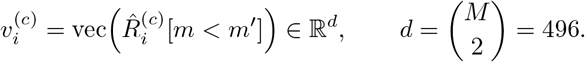

The representation is fixed-dimension across patients (it does not depend on 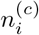) and encodes within-compartment co-ordination between biological programs rather than raw expression levels. This is the core compression step: a sparse, heterogeneous, patient-specific cells-by-genes object is replaced by a dense, comparable 496-dimensional patient vector that preserves the organization of programs within the sample.

Figure 10a shows the intermediate stage of this compression for an exemplar paired patient. Rows are cells (downsampled for display), columns are selected genes, genes are ordered by their learned module assignment, and the colored strip above the heatmap indicates module membership for each gene block. Under this ordering, coherent vertical blocks emerge that are not visible in the unordered cells-by-genes matrix. Figure 10b shows the final patient-level object 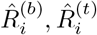 for the same patient, with rows and columns indexed by the shared module order. Each pixel encodes the dependency between two biological programs within that patient. This is the input that the encoder networks consume.

**Figure 10.**
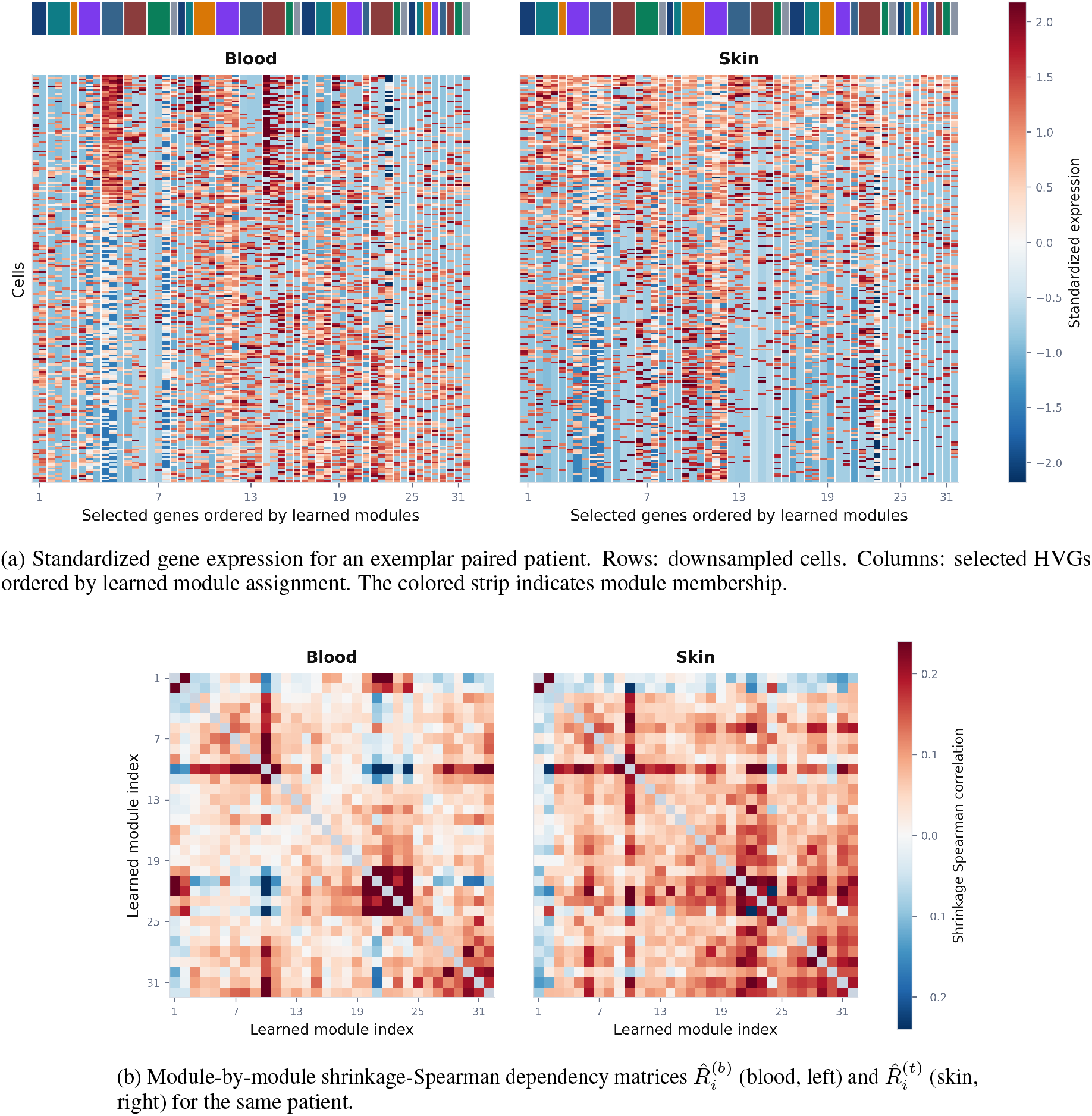
From raw expression to the patient-level dependency operator. (a) Under the learned module ordering, coherent gene blocks emerge in the standardized cells-by-genes view of an exemplar patient. (b) The second compression step summarizes each patient and compartment as a 32 × 32 dependency matrix whose strict upper triangle (*d* = 496 entries) is the fixed-length patient vector consumed by the encoder.

### 8.3 Architecture and objective

The encoders *f*_*b*_, *f*_*t*_: ℝ^496^ → ℝ^32^ are MLPs with hidden widths (256, 128), GELU activations, dropout 0.10, and layer normalization. Applied to the patient vectors they produce raw latents 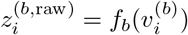 and 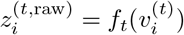; these are *ℓ*_2_-normalized to unit length to give alignment latents

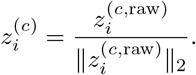

Decoders *g*_*b*_, *g*_*t*_: ℝ^32^ → ℝ^496^ mirror the encoders with hidden widths (128, 256). They are evaluated on four paths: self-reconstruction 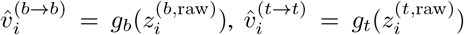, and cross-reconstruction 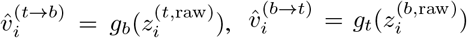. Two supervised heads *h*_*b*_, *h*_*t*_ with a single hidden layer of width 64 predict the continuous tissue-target vector 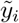 (standardized using training-fold statistics) from each compartment latent:

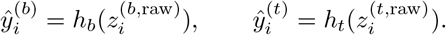

The training objective is a weighted sum of seven terms:

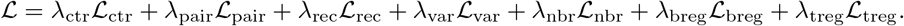

The contrastive term ℒ_ctr_ is a symmetric InfoNCE on the cross-view logit matrix *ℓ*_*ij*_ = *z*^(*b*)^ *z*^(*t*)^*/τ* with temperature *τ* = 0.20,

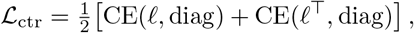

which matches each blood sample to its paired tissue sample against all other tissue samples in the batch, and vice versa. The pair cosine term enforces direct alignment of matched pairs,

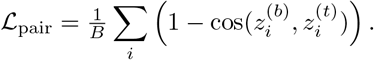

The reconstruction term uses a masked Huber loss with *δ* = 1.0, combining self- and cross-reconstruction,

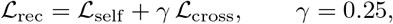

which forces the latent to encode enough structure to reconstruct each compartment’s dependency matrix from its own latent and from the other compartment’s. The variance term prevents latent collapse by penalizing low per-dimension variance across the batch,

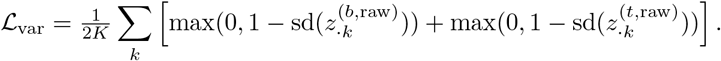

The neighborhood term preserves patient geometry from the input space to the latent space by matching off-diagonal entries of the cosine-similarity matrix,

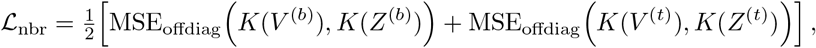

where 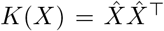 is the cosine-similarity kernel on *ℓ*_2_-normalized rows of *X*. The two supervised heads are trained with masked Huber regression on the standardized tissue target,

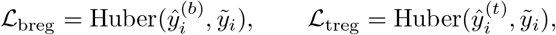

which forces the latent to be directly useful for tissue-target prediction from both sides. Loss weights are set to *λ*_ctr_ = 1.0, *λ*_pair_ = 0.25, *λ*_rec_ = 0.75, *λ*_var_ = 0.05, *λ*_nbr_ = 0.10, *λ*_breg_ = 1.0, *λ*_treg_ = 0.50.

### 8.4 Training and geometry

The quantitative evaluation uses AdamW with learning rate 10^−3^, weight decay 10^−4^, batch size 8 patients, gradient clipping at norm 1.0, and a maximum of 50 epochs. An internal validation split (20% of training patients) is used for early stopping on the blood-side regression loss with a patience of 10 epochs. Performance is reported under 5-fold patient-level cross-validation with 1 repeat. Basis construction, target standardization, model fitting, and early stopping are all performed strictly on training-fold patients; test patients contribute nothing to any of these steps.

Two training-time null families are run under the same protocol. The shuffled-pair null constructs batches 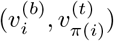 with a non-identity permutation *π*, breaking patient-level blood–tissue pairing. The target-permuted null constructs batches 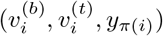, keeping the paired inputs but breaking target supervision. Simple baselines include cosine retrieval directly on the matrix-view patient vectors, ridge regression from 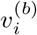 to *y*_*i*_, and partial least squares.

The projection figures in this section are descriptive rather than held-out evaluation. Because latent spaces produced by separately trained outer-fold models are not directly comparable, the projection geometry is not constructed by pooling latents across cross-validation folds. Instead, for visualization only, one internally validated fit is trained per analysis context using a single train/validation split for early stopping, and all patients are then embedded in that single coherent latent space. These geometry figures are used to interpret the learned structure of the model; the held-out evidence remains the cross-validated target-recovery analysis below.

### 8.5 Results

Figure 11 shows the learned patient geometry in the descriptive single-fit latent space. Tissue latents 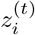 are embedded in the leading principal components of the tissue latent; blood latents 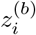 are projected into the same PCA basis. *K*-means with silhouette-selected *k* is run on the tissue latents, and points are colored by the resulting internally learned neighborhoods. Filled points are tissue embeddings; open circles are blood-only projections; faint line segments link matched blood and tissue from the same patient. The figure should be read as neighborhood-level geometry rather than exact patient retrieval: blood projections tend to fall near the matched tissue neighborhood. The matched projection remains substantially more coherent than both a shuffled-pair control and a raw-cosine baseline, supporting the interpretation that the learned geometry reflects paired cross-compartment structure rather than a trivial projection artifact.

**Figure 11.**
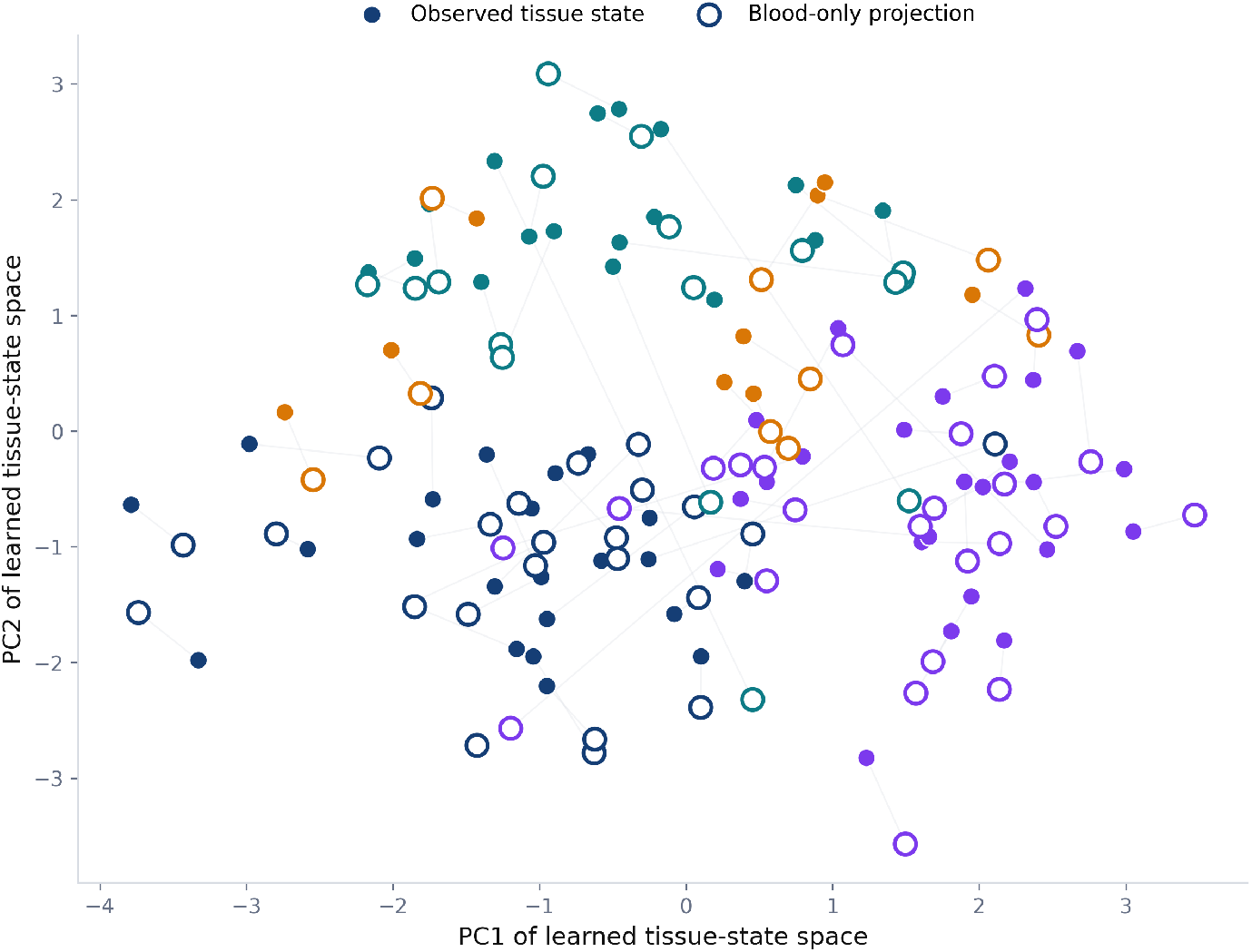
Descriptive patient geometry in the learned latent space from a single internally validated fit. Filled points: tissue embeddings 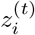. Open circles: blood-only projections 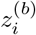 into the same tissue PCA basis. Faint line segments link matched patients. Colors denote internally learned tissue neighborhoods in the tissue latent. The figure is intended to show neighborhood-level blood-to-tissue placement, not exact patient-identity recovery.

Under the stricter held-out patient-retrieval evaluation within SSc, the exact matched tissue sample is ranked first for 13.9% of blood patients, within the top 3 for 27.9%, and within the top 5 for 47.3%. Here recall@*k* denotes the fraction of held-out blood patients for whom the true matched tissue patient appears among the top *k* ranked tissue candidates. Because each blood sample is matched against 57 possible SSc tissue patients, random top-1 retrieval would be only 1*/*57 ≈ 1.8%, so a recall@1 of 13.9% is substantially above chance even though exact identity recovery remains difficult. The mean rank of the true matched tissue patient is 5.98 and the median rank is 5.9, meaning that the correct tissue sample is typically placed around sixth out of 57 candidates rather than near the middle of the ranked list. This pattern is consistent with a model that has learned substantial patient-level cross-compartment structure, but recovers coarse tissue-state neighborhoods more reliably than exact patient identity.

Figure 12 shows the held-out quantitative result. The blood-side latent readout is evaluated on continuous tissue targets under patient-level cross-validation and compared against the best simple baseline on the same patients and folds. The comparison metric is mean held-out Spearman correlation. The gray interval is a conservative null band spanning the upper control range from the target-permuted and shuffled-pair training-time null families. On several held-out tissue targets the latent readout exceeds both the best simple baseline and the null band, indicating that paired multiview training extracts information from blood that is not captured by the same patient vectors under simpler linear readouts or under broken pairing/supervision controls. Illustrative examples include skin LC, fibroblast MYOC2 proportion, and tissue fatty-acid metabolism in the all-paired setting, and UV response up, vascular ACKR1 proportion, and mitotic spindle within SSc.

**Figure 12.**
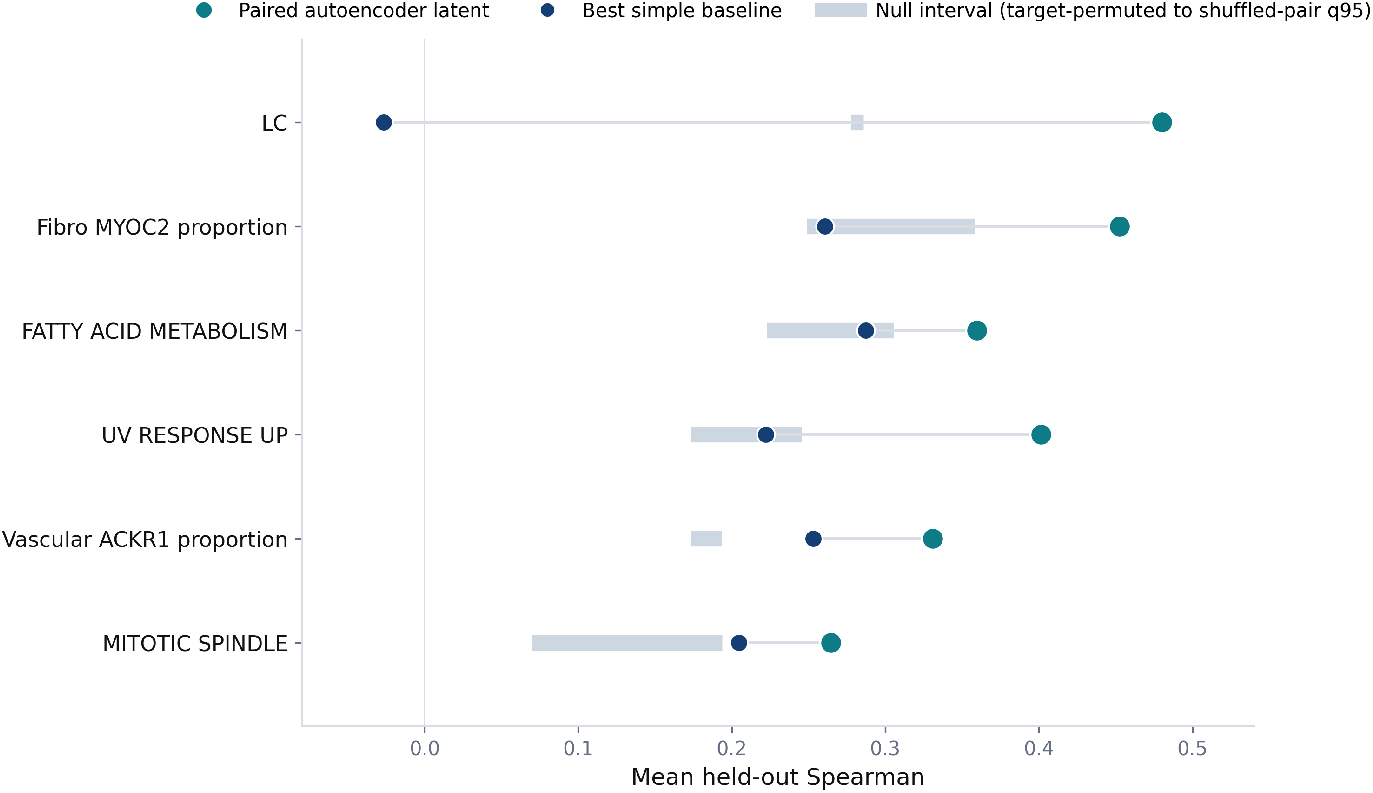
Held-out continuous tissue-target recovery from the autoencoder latent. Teal: blood-side latent readout. Blue: best simple baseline on the same matrix-view patient vectors. Grey interval: conservative null band spanning the target-permuted and shuffled-pair training controls. Points are mean held-out Spearman correlation under patient-level cross-validation.

## 9 Discussion

The paired Gur cohort contains enough cross-compartment structure to recover tissue-defined molecular states from blood at patient level. The bridge is visible in univariate cross-compartment correlations, in shared multivariate geometry over the joint pathway space, in residualized cell-type-resolution association maps, and in out-of-fold classification of tissue-defined labels from blood features alone. It extends beyond interferon into fibroblast, stromal, vascular, proliferative and metabolic tissue programs, with the strongest classical-ML recoveries on tissue-resident fibroblast subtypes whose only route into blood is through the indirect mechanism by which an active fibroblast microenvironment reshapes circulating immune populations. The same cross-compartment structure is learnable end to end: the paired autoencoder places blood-only projections into the same learned tissue-state neighborhood as the corresponding tissue samples, and blood-side readout from that latent clears two permutation null families on selected held-out tissue targets.

The evidence here is single-cohort and external replication in an independent paired dataset is the next step; until that step lands, the quantitative claims above apply to the Gur cohort. Tissue targets in sections 4 to 6 are defined by median splits on cohort-internal distributions of skin gene-set scores and cell-state proportions. Anchoring a subset of these targets against clinically validated tissue programs — for instance aligning fibroblast POSTN recovery against mRSS in an independent dcSSc cohort with paired biopsy and skin scoring — will upgrade the central claim from predicting an internal split to predicting a clinically established tissue program. The single-cell interface used across the classical analyses is a three-statistic compression per cell-type–gene-set pair; the scaling curve in section 7 is still rising at *n* = 74 under this compression alone, and richer distributional features become statistically estimable at *n* = 200–500.

The modeling framework is indication-agnostic. Autoimmune diseases share a common architecture of tissue-localized pathology driven by immune dysregulation, with circulating immune populations that traffic between blood and the affected organ. Coupling mechanisms vary by indication — synovial macrophage polarization in rheumatoid arthritis, plasma cell infiltration and complement deposition in lupus nephritis, alveolar macrophage reprogramming in SSc-associated interstitial lung disease, and epithelial–immune crosstalk in inflammatory bowel disease — and each paired cohort contributes supervision on a different region of the autoimmune tissue-state manifold. Paired scRNA-seq cohorts in other indications are either publicly available (AMP Phase II in rheumatoid arthritis, 314,000+ cells across 79 donors) or in active collection.

The deployment interface is not restricted to scRNA-seq. Plasma proteomics measures thousands of circulating proteins that reflect tissue-level processes through secretion and cell-surface shedding; the same fibroblast remodeling programs that reshape blood T-cell composition in this study also shed COMP and MMP-1 into plasma at levels that correlate with mRSS [4]. Cell-free DNA methylation profiling detects epigenetic signatures of dying cells, including tissue-resident cell types that do not circulate. Each modality is a partial view of the same latent tissue state. Training on paired scRNA-seq grounds the latent in cellular and distributional biology, and adding proteomic and cfDNA views measured on the same patients tightens the posterior on axes where transcriptomic data alone is underpowered.

SSc is the first demonstration. The broader target is a single model for autoimmune tissue state, trained on paired blood–tissue single-cell data across indications and deployed through peripheral blood. The classical and learned results presented here define the empirical geometry of such a model at *n* = 74 in one indication, and the scaling behavior indicates that its fidelity grows with cohort size.

## Notes

https://www.ncbi.nlm.nih.gov/geo/query/acc.cgi?acc=GSE195452

### Competing Interest Statement

The authors have declared no competing interest.

